# Adjusting for Variable Brain Coverage in Voxel-Based fMRI Meta-Analysis

**DOI:** 10.1101/457028

**Authors:** Jo Cutler, Joaquim Radua, Daniel Campbell-Meiklejohn

## Abstract

Meta-analyses of fMRI studies are vital to establish consistent findings across the literature. However, fMRI data are susceptible to signal dropout (i.e. incomplete brain coverage), which varies across studies and brain regions. In other words, for some brain regions, only a variable subset of the studies included in an fMRI meta-analysis have data present. These missing data can mean activations in fMRI meta-analysis are underestimated (type II errors). Here we present SPM (MATLAB) code to run a novel method of adjusting random-effects models for meta-analytic averaging of a group of studies and mixed-effects models for comparison between two groups of studies. In two separate datasets, meta-analytic effect sizes and z-scores were larger in the adjusted, compared to the unadjusted analysis. Relevantly, these changes were in regions such as the ventromedial prefrontal cortex where coverage was lowest. Limitations of the method, including issues of how to threshold the adjusted maps are discussed. Code and demonstration data for the adjusted method are available at https://doi.org/10.25377/sussex.c.4223411.

## 1. Introduction

Echo-planar images for functional magnetic resonance imaging (fMRI) are susceptible to signal dropout (Ojemann et al., 1997) leaving gaps in activation maps. The level of coverage can vary widely between individuals, scanners, and scan protocols. This presents a problem of false negatives (type II errors – wrongly concluding no effect exists) for both individual studies and for map-based fMRI meta-analyses.

The problem of missing data from lack of coverage is limited to a subset of regions, including ventromedial prefrontal cortex (vmPFC) and temporal lobes, due to factors such as nearby air and bone. False negatives are therefore localised to these regions and not uniformly distributed throughout the brain. Research on topics and tasks which rely on these regions, for example value-based decision-making in vmPFC (Levy and Glimcher, 2012), will be disproportionately affected by issues of coverage. In addition, if meta-analyses test for convergence, incomplete brain coverages may lead to incorrect p-values, because the test assumes that “false foci” are uniformly distributed across the brain (Albajes-Eizagirre and Radua, 2018).

While techniques have been developed to maximise coverage (Weiskopf et al., 2007), they are not uniformly successful or universally applied. In this report, we discuss an approach to reducing these type II errors in map-based fMRI meta-analyses.

Using meta-analysis techniques on neuroimaging data is vital to establish consistent neural correlates across studies (Müller et al., 2018; Wager et al., 2007, 2009). Several tools are available. One technique for meta-analysis is Anisotropic Effect Size Signed Differential Mapping software (AES-SDM, Radua et al., 2014) which combines coordinate-based meta-analysis with unthresholded maps (Radua and Mataix-Cols, 2012) to reduce assumptions of the spatial extent of activations.

Dropout in where signal is present can mean activations in fMRI meta-analysis are missed or underestimated. This will be, at least in part, due to voxels where no effect size was measured, being attributed the same variance estimates as voxels where effect sizes were measured. Here we present code which runs a novel method of adjusting both random and mixed-effects models, for meta-analytic averaging across a single group or comparison between two groups of studies respectively. The code adjusts each type of variance (within-study & between-study) in the models used in AES-SDM, which are usually assigned to every voxel, so only voxels where data was recorded are included.

## 2. Materials and methods

### 2.1. Data selection

This technique to account for variable coverage was developed as part of an fMRI meta-analysis on prosocial behaviour so a detailed description of selection methods is provided elsewhere (Cutler and Campbell-Meiklejohn, 2019). Briefly, meta-analyses were calculated for each of two groups of decision type, “altruistic” and “strategic”, using random-effects models and these groups were compared with a mixed-effects model. The altruistic group contained 18 maps and 3 coordinates sets (n = 21, 557 participants) while the strategic group had 10 maps and 5 coordinates sets (n = 15, 593 participants). Due to different control conditions across studies, the adjusted analysis was only run on studies which contrast altruistic (n = 12) or strategic (n = 12) with selfish decisions.

To establish the wider relevance of the method, we conducted a second meta-analysis using data which researchers have made available through NeuroVault (Gorgolewski et al., 2015). It is vital to stress that this is in no way a comprehensive meta-analysis of any tasks and it is unlikely that a genuine meta-analysis would group these maps together. These maps were simply used as their CC0 license enables sharing as a demonstration set with the code (available at https://doi.org/10.25377/sussex.c.4223411).

Searches on NeuroVault were conducted for “choice” and “deci*” (for decision, decide etc.). Maps were selected if they had data from any decision task in the scanner with a contrast to no decision or a decision which varied on a parameter, for example complexity. This crude selection technique resulted in 18 maps (see Table 1).

**Table 1.** Details of the studies with data available from NeuroVault included in the second meta-analysis

| First author | Year | NeuroVault collection name | Map used | Map type | Sample size |
| --- | --- | --- | --- | --- | --- |
| Park | 2017 | Integration of individual and social information for decision-making in groups of different sizes | Brain regions for changing judgments to conform to the group's | T | 23 |
| Chang | 2011 | Great expectations: neural computations underlying the use of social norms in decision-making | Offer3_BetweenExpect | Z | 17 |
| Bang | 2018 | Distinct encoding of decision confidence in human medial prefrontal cortex | GLM1: Group-level main effect of coherence (un-masked) | T | 32 |
| Fleming | 2018 | Neural mediators of changes of mind about perceptual decisions | Parametric effect of reported confidence | T | 22 |
| Gonzalez Alam | 2018 | Meaningful inhibition: Exploring the role of meaning and modality in response inhibition | Figure 2. Semantic and Perceptual Inhibition Overlap | Z | 27 |
| Tom | 2007 | The Neural Basis of Loss Aversion in Decision-Making Under Risk | task001_cope03_parametric gain | T | 16 |
| Waskom | 2016 | Adaptive Engagement of Cognitive Control in Context-Dependent Decision Making | Context prediction error | Z | 15 |
| Op de Macks | 2018 | Supplement: Neural correlates of adolescent decision-making and outcome processing during the Yellow Light Game | Stop > Go (N=78, 11-17 years-old, community sample) | T | 78 |
| Suzuki | 2015 | Neural Mechanisms Underlying Human Consensus Decision-Making | Fig.4A (Main + Control) | T | 20 |
| Rahnev | 2016 | Causal evidence for frontal cortex organization for perceptual decision making | Stimulus/perceptual judgment epoch | T | 17 |
| Li | 2017 | Reason's Enemy Is Not Emotion: Engagement of Cognitive Control Networks Explains Biases in Gain/Loss Framing | Frame-Consistent>Framing-Inconsistent, unthresholded | Z | 143 |
| Aridan | PreP | Neural correlates of effort-based valuation with prospective choices | effort_gain_pos | Z | 40 |
| Cho | 2016 | Reward Sensitivity Enhances Ventrolateral Prefrontal Cortex Activation during Free Choice | Figure2 Free vs Forced TEPS unthresh | Z | 33 |
| Kameda | 2016 | Rawlsian maximin rule operates as a common cognitive anchor in distributive justice and risky decisions | Activity of the right temporo-parietal junction (RTPJ) during decision making. | T | 30 |
| van der Laan | 2014 | Sweet lies: neural, visual, and behavioral measures reveal a lack of self-control conflict during food choice in weight-concerned women | Choice periods SC vs NSC | T | 20 |
| Hunt | 2014 | Hierarchical competitions subserving multi-attribute choice | Integrated value competition | Z | 19 |
| Korn | 2018 | Heuristic and optimal policy computations in the human brain during sequential decision-making | GLM2: Participants' choices (binary parametric modulator: foraging versus waiting) | T | 28 |
| Fujiwara | 2018 | Ventrolateral Prefrontal Cortex Updates Chosen Value According to Choice Set Size | spmT 0001 | T | 18 |

Coverage was investigated by binarising each map, after registration to a common template, based on whether there was signal in each voxel and summing these images to create coverage maps (Figure 2). Both the dataset on prosocial decisions and the NeuroVault dataset on decisions showed decreased coverage around the periphery, particularly in vmPFC.

### 2.2. Combined image and coordinate meta-analysis

Using maps in fMRI meta-analysis has a number of benefits including enhanced sensitivity and detection of consistent but subthreshold effects. When maps are unavailable, AES-SDM recreates estimated maps from coordinates and their effect sizes using an anisotropic kernel. If obtainable, peaks can be entered in both directions of the contrast.

Statistics other than T are transformed before all maps, including those recreated from coordinates, are aligned to a common template. The software then implements a permutation-based analysis. The recommendation is 50 permutations which creates 50 randomisations with the same number of foci as the map of interest. These preprocessing steps result in recreated NIfTI maps of effect sizes and within-study variance for each study. These maps are used in the both the original, unadjusted method by the software and the adjusted technique described here. Adjusted analysis uses custom scripts in SPM12 (Statistical Parametric Mapping, http://www.fil.ion.ucl.ac.uk/spm) which are available under an MIT license from https://doi.org/10.25377/sussex.c.4223411 and github.com/jocutler/adjusting-dropout-fMRI-meta.

### 2.3. Random-effects model

#### 2.3.1. Unadjusted model

One widespread use of meta-analysis is to calculate mean effect sizes across studies. A common method, and the method used in AES-SDM, is a random-effects model. In the model, AES-SDM weights each study by the inverse of the total (within-study and between-study) variance. The between-study variance, τ^2^, is obtained by the DerSimonian-Laird estimator (Dersimonian and Laird, 1986) as:

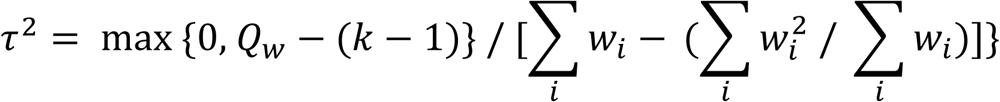

Where w_i_ is a weighting calculated as the inverse of the ith study’s within-study variance, k is the number of studies and Qw is calculated as:

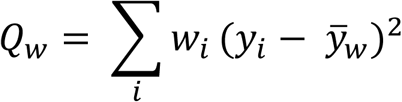

Where y_i_ is the ith study effect size estimate, wi is a weighting calculated as the inverse of the ith study’s within-study variance and 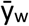 is the weighted estimate of the overall effect size calculated as:

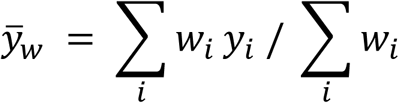

In simplified terms, used in the code:

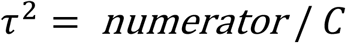

Where:

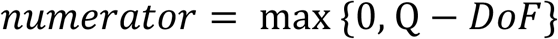

With Q calculated as above, degrees of freedom (DoF) the number of studies -1 and:

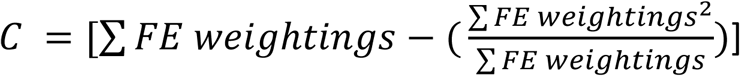

Where FE weightings are the inverse of the studies’ within-study variance. These are referred to as fixed-effects (FE) weightings as they are the ones used for a fixed-effects model, which just takes into account within-study (not between-study) variance.

In practice, these equations demonstrate that the between-study variance (τ^2^) depends on the sum of the weightings (w_i_) which are the inverse of the within-study variance.

#### 2.3.2. Adjusting random-effects models

In fMRI meta-analysis, the within-study variance is a single number which is applied as the variance across every voxel in the brain within the mask of interest. The effect size for that voxel is the transformed effect size created during preprocessing. However, if signal dropout has occurred, the effect size is zero. This means that voxels with no recorded signal are attributed variance but no effect size. When the FE weightings and effect sizes are each summed during calculation of the between-study variance, these voxels are contributing to the total variance without contributing an effect size. This could underestimate meta-analytic effect size due to inflated variance.

To account for this, calculations for the meta-analysis can be adjusted so only studies where data was recorded contribute weightings to the calculation of τ^2^. This can be done either at the single-voxel level with a spreadsheet (Figure 1) or across the whole brain, voxel-by-voxel, by masking variance maps with their coverage. The DoF value is also adjusted to be the number of studies with data – 1.

**Figure 1.**
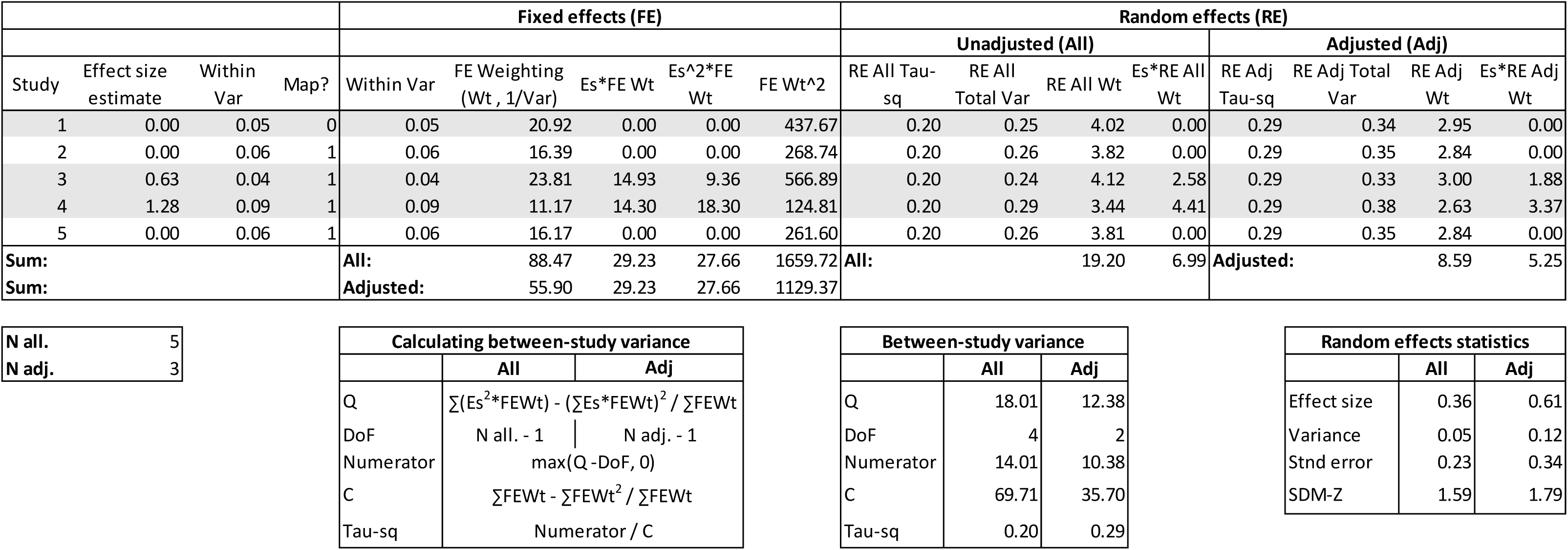
Demonstration of an adjusted random-effects model on a single voxel for five studies. Shaded rows (studies 1, 3 & 4) are those which are still taken into account in the adjusted analysis. “All” refers to the unadjusted analysis with all studies. Study 1 is from coordinates so this is included despite having an effect-size estimate of zero as this could be meaningful (see section 2.3.2 for details and exception to this rule). Studies 3 and 4 are maps with non-zero effect-size estimates in the voxel of interest, so are included. Studies 2 and 5 are maps with zero effect-size estimates, suggesting missing data due to signal dropout, so these are not included in the adjusted analysis. The fixed-effects section is purely used to calculate the between-study variance (τ^2^: tau-sq) with the calculations for this shown in the box “calculating between-study variance”. Although τ^2^ increases from 0.20 to 0.29 in the adjusted analysis, likely linked to having less studies, the meta-analytic effect size (g) increases substantially from 0.36 to 0.61 as the sum of the RE weightings (which the sum of the weighted effect sizes is divided by) decreases from 19.20 to 8.59. An interactive spreadsheet in this format can be downloaded from https://doi.org/10.25377/sussex.c.4223411 to run this analysis on any voxel for any set of studies.

It is important to note that maps recreated from coordinates should *not* be adjusted *unless* the coverage is known. If the coverage is unknown, effect sizes with values of zero do not necessarily imply lack of signal and could meaningfully demonstrate the voxel is too far from any peaks to be attributed effect size. Of course, the maps which these coordinates were generated from could also suffer from signal dropout but this cannot be confirmed. This is another reason to obtain maps wherever possible.

If the coverage is known, for example if the paper states the cerebellum was not analysed, a coverage map reflecting this could be created and entered into the analysis as the mask for that study. This was not done for any of the studies in the current analyses.

Once τ^2^ has been calculated as a single number for the between-study variance, it is added to the within-study variance for each study to provide total variance. The inverse of this total variance provides the random-effects (RE) weightings for each study, by which the effect-size estimates are multiplied.

The issue of variable coverage affects results again at this stage as the overall meta-analytic effect size (Hedges’ g) is calculated by the sum of the weighted effect sizes divided by the RE weightings summed:

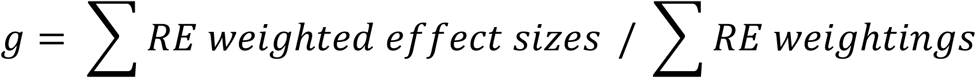

The meta-analytic variance map, used to calculate standard error and z-scores, is calculated as the inverse of the RE weightings summed:

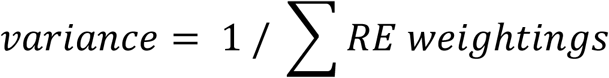

Again, voxels with no effect-size estimate due to missing data will contribute zero to the effect sizes but increase the sum of the RE weightings in both of these calculations. A greater value of summed weightings leads to underestimation of g and overestimation of variance.

The same principle can be applied here as with the FE weightings, where studies are only included in the weightings sum if they have an effect size present.

### 2.4. **Mixed-effects model**

#### 2.4.1. Unadjusted model

In addition to calculating the mean effect size for a group of studies, meta-analysis can calculate the difference between two groups using a mixed-effects model. The calculations and method of adjusting are similar to random-effects models, except the DoF equals the total number of studies across groups -2, the calculation of Q is:

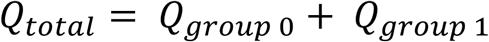

And the calculation of C is:

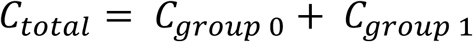

Where Q and C for each group separately are calculated as above. Groups are referred to as 0 and 1 to match dummy coding in AES-SDM.

Calculating τ^2^ follows the same process as above to produce a single number for the between-study variance across all the studies in both groups. This is added to the within-study variance and the inverse of this total variance provides the study’s mixed-effects (ME) weighting.

The meta-analytic effect size (Hedges’ g) is then calculated with the formula shown above for *each group of studies separately* - the sum of ME weighted effect sizes divided by the sum of ME weightings for that group. The meta-analytic variance is also calculated as above for each group of studies separately: the inverse of the ME weightings sum for that group.

The Hedges’ g effect-size map for the difference between groups is the calculated by *subtracting* the two separate effect-size maps:

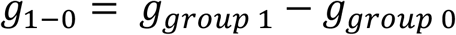

The meta-analytic variance for the difference between groups is calculated by *summing* the two separate variance estimates:

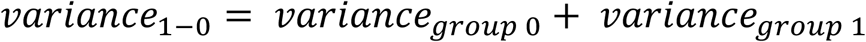

#### 2.4.2. Adjusted model

As with random-effects models, voxels with zero effect sizes due to dropout are attributed within-study variance and so increase the sums of FE and ME weightings. This is likely to underestimate average effect sizes. The same adjustments can avoid this problem in a mixed-effects model by excluding weightings of voxels where no effect size is present, unless the map was recreated from peaks with unknown coverage. This adjustment can again be done for a single voxel or across the whole brain. The DoF becomes the total number of studies, across both groups, which contribute weightings – 2.

### 2.5. Z-maps and thresholding

Once the effect-size and variance maps have been adjusted, maps of standard error and z-scores can be produced. As the input came from permutations in AES-SDM, z-scores are “SDM-Z” because they do not follow a normal distribution.

Standard error (SE) is the square root of the meta-analytic variance, either for a single group or the difference between groups (shown):

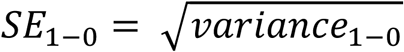

SDM-Z is the effect size (Hedges’ g) divided by the standard error, either for a single group or the difference between groups (shown):

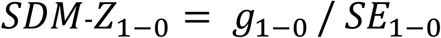

Thresholding in AES-SDM uses a voxel-level threshold of *p*<0.005 which approximates *p*<0.05 corrected and balances specificity and sensitivity (Radua et al., 2012). However, in SDM-Z maps from the adjusted method, voxels have differing DoF meaning thresholding is not straightforward.

In the prosocial decisions meta-analysis (Cutler and Campbell-Meiklejohn, 2019), maps were thresholded with SDM-Z > 2.3. This was chosen as a common value for thresholding, close to the average of the critical SDM-Z values generated in the original, unadjusted analyses (with all studies) and AES-SDM analyses run with the 50% of maps with the best coverage. Here, we apply the same threshold to the NeuroVault data. We recognise this is not a perfect solution but it provides continuity and this analysis is not meaningful, regardless of thresholding method, other than for demonstrating the impact of the adjustment.

## 3. Results

In both random and mixed-effects analyses, in both datasets, the adjusted method increased effect sizes across the lower vmPFC where coverage was worst (Figure 2). Activations based on SDM-Z > 2.3 uncorrected were larger in the adjusted than the unadjusted analysis.

**Figure 2.**
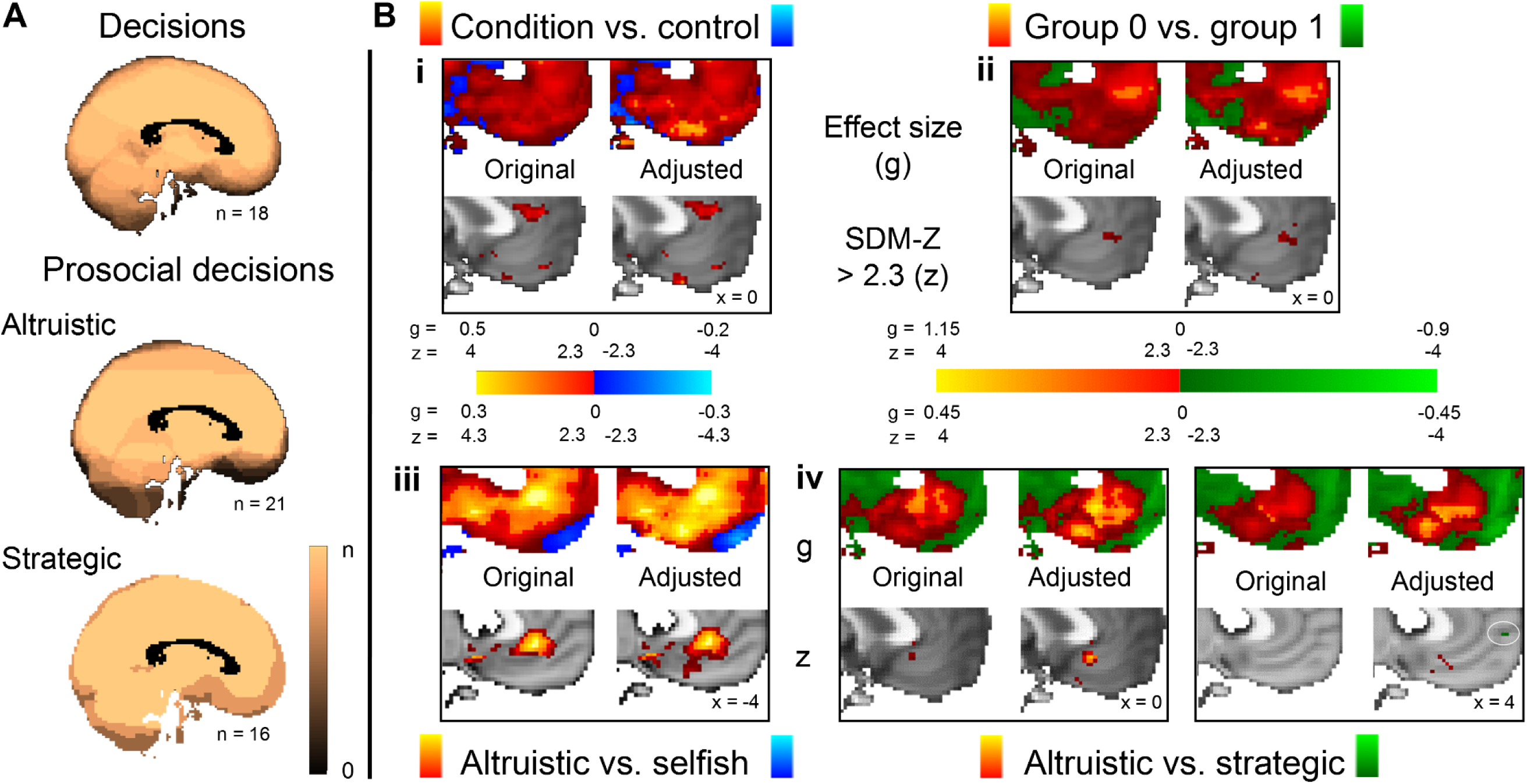
Coverage maps and results. **(A)** Coverage maps showing the number of studies with data in each region, x = 0, n = the number of studies available and the maximum possible coverage. **(B)** Increased effect sizes (Hedges’ g; top rows) and larger regions of SDM-Z > 2.3 (bottom rows) in the adjusted analysis accounting for vmPFC signal dropout, compared to the unadjusted analysis with all studies. Decisions (data from NeuroVault; n = 18) **(i)** average (random-effects) and **(ii)** comparison (mixed-effects). Prosocial decisions **(iii)** altruistic average (n = 12; random-effects) **(iv)** comparison between altruistic and strategic decisions (n = 24; mixed-effects).

### 3.1. Prosocial decisions

In the prosocial decisions data, effect sizes and the size of SDM-Z > 3 activations increased in the adjusted analysis for altruistic vs. selfish (Figure 2b iii) and altruistic vs. strategic (Figure 2b iv). For altruistic > strategic in posterior vmPFC and strategic > altruistic in anterior vmPFC, some activations were shown only in the adjusted analysis. This dissociation fits with findings of a posterior to anterior vmPFC axis differentiating altruistic from strategic decisions (Cutler and Campbell-Meiklejohn, 2019).

### 3.2. Decision-making (NeuroVault data)

Results from the second dataset from NeuroVault on decision-making further support the use of the adjusted method to account for coverage. Again, it is vital to stress that this analysis purely provides a second application of the adjustments for signal dropout and results are not a comprehensive meta-analysis or a meaningful group.

Effect sizes across vmPFC increased in the adjusted analysis in both the random-effects model on all 18 studies’ decision-making condition vs. control (Figure 2b i) and the mixed-effects model comparing 2 randomly-allocated groups of 9 studies each (Figure 2b ii). Similarly, regions where SDM-Z > 2.3 were larger or only present in the adjusted analysis.

## 4. Discussion

This paper presents a novel method of adjusting voxel-based fMRI meta-analysis technique AES-SDM (Radua et al., 2014) to account for variable brain coverage between studies. Increased effect sizes and larger regions with substantial z-scores were found in the adjusted, compared to the unadjusted analysis, using two datasets.

Our results suggest that regions that suffer from signal dropout, such as vmPFC, can show false negatives if coverage is not accounted for. The role of these regions may be underestimated or overlooked, preventing a complete understanding of their function. In meta-analyses, lack of coverage in some studies may obscure an activation despite high consistency in the studies with data in that region.

The uneven spatial distribution of signal dropout may also increase false positive results in the rest of the brain. Specifically, the test for convergence used in current coordinate-based meta-analyses assumes a uniform distribution of false positive study peaks (Albajes-Eizagirre and Radua, 2018), but this assumptions is incorrect if some brain regions have no data in some studies. This additional problem should not happen in the upcoming version of SDM, which no longer conducts tests for convergence.

To overcome these issues, our method adjusts random and mixed-effects meta-analysis models to only include variance, at each calculation stage, from voxels with study-level effect-size data. This means the number of contributing studies ranges between one and the total number of studies. The voxel degrees of freedom will be the number of contributing studies -1 (random-effects model) or -2 (mixed-effects model). For some voxels with only a few studies contributing, single studies could dominate and prevent the benefits of meta-analysis from being realised. In comparisons between two groups, differences between the sizes of each group could be driven by coverage.

Perhaps the biggest challenge from varying degrees of freedom across voxels is thresholding the adjusted meta-analytic map. In the prosocial decisions meta-analysis, SDM-Z > 2.3 was applied to the adjusted and unadjusted analysis as a comparison (Cutler and Campbell-Meiklejohn, 2019). Although this is liberal for z-scores which follow a normal distribution, SDM-Z scores do not follow a normal distribution (Radua et al., 2012) 2.3 was close to the threshold SDM-Z score generated in the unadjusted analysis with all studies and the half with the best coverage. Here, we also threshold the second meta-analysis with SDM-Z > 2.3 for continuity.

This is not a perfect solution and this adjusted method is perhaps best used alongside unadjusted analyses thresholded in a more appropriate way. The usefulness of a simple SDM-Z threshold may be limited to visual comparisons between unadjusted and adjusted maps. Comparing adjusted and unadjusted effect-size maps (Hedges’ g) can also reveal the impact of coverage deficits.

Evidence for the importance of adjusting calculations is presented using two datasets. One is a comprehensive meta-analysis of prosocial decisions. The second group is studies with data on NeuroVault for decision tasks. Decision-making was chosen as it activates vmPFC (Levy and Glimcher, 2012), a region prone to reduced coverage. This data was purely used as the availability enables sharing to demonstrate the code and results are not be meaningful in any other way. That the NeuroVault data did not show large vmPFC activations is likely due to variety in tasks, which would not be grouped together in a genuine meta-analysis.

## 5. Conclusion

Using meta-analysis techniques on fMRI data establishes consistent findings across samples, scanning sites and tasks, providing many benefits and overcoming issues with single studies. Including statistical maps enhances some of these benefits and methods like AES-SDM, which combine maps with coordinates, increase the chances of including a study. However, fMRI meta-analyses are prone to false negatives if lacking coverage leaves missing data in study-level statistical maps. Several key regions, including vmPFC and temporal areas suffer from signal dropout, meaning false negatives are unevenly distributed throughout the brain.

To account for and overcome these issues, we present a novel method of adjusting calculations for random and mixed-effects models for meta-analytic group averages and comparisons respectively. By adjusting models to only include variance for voxels with data present in the study’s effect-size map, we demonstrate increased meta-analytic effect sizes in regions with the worst coverage. This highlights that failing to account for coverage underestimates effect sizes or may miss activations altogether.

## Acknowledgements

Thank you to Anton Albajes-Eizagirre who provided feedback during the development of the method and co-developed AES-SDM. We also thank all of the authors who provided or looked for data to contribute to the prosocial decisions meta-analysis, the authors who shared their data via NeuroVault and the creators of that platform for enabling and encouraging data sharing.

## Author contributions

JC and DC created the concept of the method which relies on the output of software developed by JR. JC wrote the code and the paper with contributions to both from DC and JR. All authors contributed to manuscript revision, read and approved the submitted version.

## References

Albajes-Eizagirre, A., and Radua, J. (2018). What do results from coordinate-based meta-analyses tell us? Neuroimage 176, 550–553. doi:10.1016/J.NEUROIMAGE.2018.04.065.

Button, K. S., Ioannidis, J. P. a, Mokrysz, C., Nosek, B. a, Flint, J., Robinson, E. S. J., et al. (2013). Power failure: Why small sample size undermines the reliability of neuroscience. Nat. Rev. Neurosci. 14, 365–76. doi:10.1038/nrn3475.

Cremers, H. R., Wager, T. D., and Yarkoni, T. (2017). The relation between statistical power and inference in fMRI. PLoS One 12, e0184923. doi:10.1371/journal.pone.0184923.

Cutler, J., and Campbell-Meiklejohn, D. (2019). A comparative fMRI meta-analysis of altruistic and strategic decisions to give. Neuroimage 184, 227–241. doi:10.1016/J.NEUROIMAGE.2018.09.009.

Dersimonian, R., and Laird, N. (1986). Meta-Analysis in Clinical Trials. Stat. Med. 188, 177–188. doi:10.1016/0197-2456(86)90046-2.

Eklund, A., Nichols, T. E., and Knutsson, H. (2016). Cluster failure: Why fMRI inferences for spatial extent have inflated false-positive rates. Proc. Natl. Acad. Sci. U. S. A. 113, 7900–5. doi:10.1073/pnas.1602413113.

Gorgolewski, K. J., Varoquaux, G., Rivera, G., Schwarz, Y., Ghosh, S. S., Maumet, C., et al. (2015). NeuroVault.org: a web-based repository for collecting and sharing unthresholded statistical maps of the human brain. Front. Neuroinform. 9, 1–9. doi:10.3389/fninf.2015.00008.

Levy, D. J., and Glimcher, P. W. (2012). The root of all value: A neural common currency for choice. Curr. Opin. Neurobiol. 22, 1027–1038. doi:10.1016/j.conb.2012.06.001.

Müller, V. I., Cieslik, E. C., Laird, A. R., Fox, P. T., Radua, J., Mataix-Cols, D., et al. (2018). Ten simple rules for neuroimaging meta-analysis. Neurosci. Biobehav. Rev. 84, 151–161. doi:10.1016/j.neubiorev.2017.11.012.

Ojemann, J. G., Akbudak, E., Snyder, A. Z., McKinstry, R. C., Raichle, M. E., and Conturo, T. E. (1997). Anatomic localization and quantitative analysis of gradient refocused echo-planar fMRI susceptibility artifacts. Neuroimage 6, 156–167. doi:10.1006/nimg.1997.0289.

Radua, J., and Mataix-Cols, D. (2012). Meta-analytic methods for neuroimaging data explained. Biol. Mood Anxiety Disord. 2, 6. doi:10.1186/2045-5380-2-6.

Radua, J., Mataix-Cols, D., Phillips, M. L., El-Hage, W., Kronhaus, D. M., Cardoner, N., et al. (2012). A new meta-analytic method for neuroimaging studies that combines reported peak coordinates and statistical parametric maps. Eur. Psychiatry 27, 605–611. doi:10.1016/j.eurpsy.2011.04.001.

Radua, J., Rubia, K., Canales-Rodríguez, E. J., Pomarol-Clotet, E., Fusar-Poli, P., and Mataix-Cols, D. (2014). Anisotropic kernels for coordinate-based meta-analyses of neuroimaging studies. Front. Psychiatry 5, 1–8. doi:10.3389/fpsyt.2014.00013.

Wager, T. D., Lindquist, M. A., Nichols, T. E., Kober, H., and Van Snellenberg, J. X. (2009). Evaluating the consistency and specificity of neuroimaging data using meta-analysis. Neuroimage 45, S210–S221. doi:10.1016/j.neuroimage.2008.10.061.

Wager, T. D., Lindquist, M., and Kaplan, L. (2007). Meta-analysis of functional neuroimaging data: Current and future directions. Soc. Cogn. Affect. Neurosci. 2, 150–158. doi:10.1093/scan/nsm015.

Weiskopf, N., Hutton, C., Josephs, O., Turner, R., and Deichmann, R. (2007). Optimized EPI for fMRI studies of the orbitofrontal cortex: Compensation of susceptibility-induced gradients in the readout direction. Magn. Reson. Mater. Physics, Biol. Med. 20, 39–49. doi:10.1007/s10334-006-0067-6.

## References for NeuroVault datasets

Aridan, N., Malecek, N. J., Poldrack, R. A., & Schonberg, T. (2018). Neural correlates of effort-based valuation with prospective choices. BioRxiv, 26. http://doi.org/10.1101/357327

Bang, D., & Fleming, S. M. (2018). Distinct encoding of decision confidence in human medial prefrontal cortex. Proceedings of the National Academy of Sciences, 115(23), 6082–6087. http://doi.org/10.1073/pnas.1800795115

Chang, L. J., & Sanfey, A. G. (2013). Great expectations: Neural computations underlying the use of social norms in decision-making. Social Cognitive and Affective Neuroscience, 8(3), 277–284. http://doi.org/10.1093/scan/nsr094

Cho, C., Smith, D. V., & Delgado, M. R. (2016). Reward sensitivity enhances ventrolateral prefrontal cortex activation during free choice. Frontiers in Neuroscience, 10(NOV), 529. http://doi.org/10.3389/fnins.2016.00529

Fleming, S. M., Van Der Putten, E. J., & Daw, N. D. (2018). Neural mediators of changes of mind about perceptual decisions. Nature Neuroscience, 21(4), 617–624. http://doi.org/10.1038/s41593-018-0104-6

Fujiwara, J., Usui, N., Eifuku, S., Iijima, T., Taira, M., Tsutsui, K. I., & Tobler, P. N. (2018). Ventrolateral prefrontal cortex updates chosen value according to choice set size. Journal of Cognitive Neuroscience, 30(3), 307–318. http://doi.org/10.1162/jocn_a_01207

Gonzalez Alam, T., Murphy, C., Smallwood, J., & Jefferies, E. (2018). Meaningful inhibition: Exploring the role of meaning and modality in response inhibition. NeuroImage, 181, 108–119. http://doi.org/10.1016/j.neuroimage.2018.06.074

Hunt, L. T., Dolan, R. J., & Behrens, T. E. J. (2014). Hierarchical competitions subserving multi-attribute choice. Nature Neuroscience, 17(11), 1613–1622. http://doi.org/10.1038/nn.3836

Kameda, T., Inukai, K., Higuchi, S., Ogawa, A., Kim, H., Matsuda, T., & Sakagami, M. (2016). Rawlsian maximin rule operates as a common cognitive anchor in distributive justice and risky decisions. Proceedings of the National Academy of Sciences, 113(42), 11817–11822. http://doi.org/10.1073/pnas.1602641113

Korn, C. W., & Bach, D. R. (2018). Heuristic and optimal policy computations in the human brain during sequential decision-making. Nature Communications, 9(1), 325. http://doi.org/10.1038/s41467-017-02750-3

Li, R., Smith, D. V., Clithero, J. A., Venkatraman, V., Carter, R. M., & Huettel, S. A. (2017). Reason’s Enemy Is Not Emotion: Engagement of Cognitive Control Networks Explains Biases in Gain/Loss Framing. The Journal of Neuroscience, 37(13), 3588–3598. http://doi.org/10.1523/JNEUROSCI.3486-16.2017

Op de Macks, Z. A., Flannery, J. E., Peake, S. J., Flournoy, J. C., Mobasser, A., Alberti, S. L., … Pfeifer, J. H. (2018). Novel insights from the Yellow Light Game: Safe and risky decisions differentially impact adolescent outcome-related brain function. NeuroImage, 181, 568–581. http://doi.org/10.1016/j.neuroimage.2018.06.058

Park, S. A., Goïame, S., O’Connor, D. A., & Dreher, J.-C. (2017). Integration of individual and social information for decision-making in groups of different sizes. PLOS Biology, 15(6), e2001958. http://doi.org/10.1371/journal.pbio.2001958

Rahnev, D., Nee, D. E., Riddle, J., Larson, A. S., & D’Esposito, M. (2016). Causal evidence for frontal cortex organization for perceptual decision making. Proceedings of the National Academy of Sciences, 113(21), 6059–6064. http://doi.org/10.1073/pnas.1522551113

Suzuki, S., Adachi, R., Dunne, S., Bossaerts, P., & O’Doherty, J. P. (2015). Neural mechanisms underlying human consensus decision-making. Neuron, 86(2), 591–602. http://doi.org/10.1016/j.neuron.2015.03.019

Tom, S. M., Fox, C. R., Trepel, C., & Poldrack, R. A. (2007). The neural basis of loss aversion in decision-making under risk. Science, 315(5811), 515–518. http://doi.org/10.1126/science.1134239

van der Laan, L. N., de Ridder, D. T. D., Charbonnier, L., Viergever, M. A., & Smeets, P. A. M. (2014). Sweet lies: neural, visual, and behavioral measures reveal a lack of self-control conflict during food choice in weight-concerned women. Frontiers in Behavioral Neuroscience, 8, 184. http://doi.org/10.3389/fnbeh.2014.00184

Waskom, M. L., Frank, M. C., & Wagner, A. D. (2017). Adaptive Engagement of Cognitive Control in Context-Dependent Decision Making. Cerebral Cortex, 27(2), 1270–1284. http://doi.org/10.1093/cercor/bhv333

